# Ensemble Machine Learning Approaches Predict Survival in Lower-Grade Glioma Based on Glycosphingolipid Gene Expression and Metabolic Modelling

**DOI:** 10.64898/2026.01.21.700788

**Authors:** Jack W. J. Welland, Janet E. Deane

**Affiliations:** Cambridge Institute for Medical Research, Department of Clinical Neuroscience, University of Cambridge, UK

## Abstract

Glycosphingolipids (GSLs) are essential components of biological membranes with important roles in cell signalling. Disrupted GSL metabolism is associated with malignancy across a range of cancers, with different GSLs implicated in distinct tumours. GSLs have potential mechanistic roles in cancer; however, their functions in Lower Grade Gliomas (LGGs) remain poorly understood. We present ensemble machine learning approaches using transcriptomic data from LGG, combined with GSL-specific metabolic simulations, to predict survival outcomes. The ensemble approach demonstrates effective risk stratification for LGG patients based on GSL gene expression. Pathway analysis of model-derived risk groups highlighted potential association of GSLs with cell motility, division and Wnt signalling in LGG pathology. Given the strong performance of machine learning approaches to predict outcomes and that GSLs are shed into the tumour microenvironment, GSL-based diagnostics and prognostics may prove clinically beneficial. A Python package enabling GSL-specific metabolic modelling and risk prediction from RNA-seq data is provided.

## Introduction

Lower grade gliomas (LGGs) are a type of primary brain tumour that can develop in both children and adults and can severely impact neurological function[1]. LGGs encompass World Health Organisation (WHO) grade II or grade III astrocytomas or oligodendrogliomas[2,3]. Glioblastoma Multiforme (GBM), a WHO grade IV glioma, is the most invasive malignant tumour of the central nervous system with <45% of patients surviving one year beyond diagnosis[4]. LGGs and GBM possess heterogenous morphological and molecular profiles and the transition of LGGs to GBM is a well-recognised but complex process involving genetic, epigenetic and microenvironmental changes[5,6]. This heterogeneity represents a severe challenge for precise diagnosis and treatment[4,7].

Early progression from LGG to GBM is driven by epigenetic changes including hypermethylation of MGMT (methylguanine-DNA methyltransferase) inhibiting DNA damage repair, and silencing of tumour suppressors, such as CDKN2A[6,8,9]. Further amplification of these and other pathways such as growth factor signalling as well as additional genetic changes including mutation of TP53 drive aggressive tumour growth[7,10,11]. Once transformation occurs, survival drops significantly[12]. Therefore, detailed molecular profiling and identification of key markers is critical for improved diagnosis and prognosis[13].

Glycosphingolipids (GSLs) are a family of bioactive, glycosylated lipids with known roles in brain development and brain cell function[14]. GSLs are also aberrantly expressed in several cancers[15]. GSLs are amphipathic molecules composed of a ceramide backbone that is embedded in the membrane and glycosylated headgroups with large structural diversity ranging from single sugars to complex branched glycan chains (**Fig.1A**). GSLs are enriched in the outer leaflet of the plasma membrane where they engage in cell-cell interactions, influence cell signalling and modulate immune responses[16–18]. In cancer, changes to GSL profiles contribute to cell proliferation, invasion and epithelial-to-mesenchymal transition promoting metastasis[19–21]. GSLs can regulate signal transduction by interacting with growth factor receptors influencing cell growth and apoptosis[15]. GSLs are known tumour-associated antigens (TAAs) and are used as diagnostic markers in several cancers[20]. In the context of gliomas, it has been previously shown that the key (neo)lacto-series gatekeeper enzyme B3GNT5 is associated with poor prognosis in GBM, with gene expression significantly associated with glioma stem cells and B3GNT5 knockdown resulting in decreased neurosphere formation[22]. Interestingly, similar behaviour has been observed in prostate cancer, where again B3GNT5 knockdown resulted in decreased cancer stem cell sphere formation[23].

**Figure 1.**
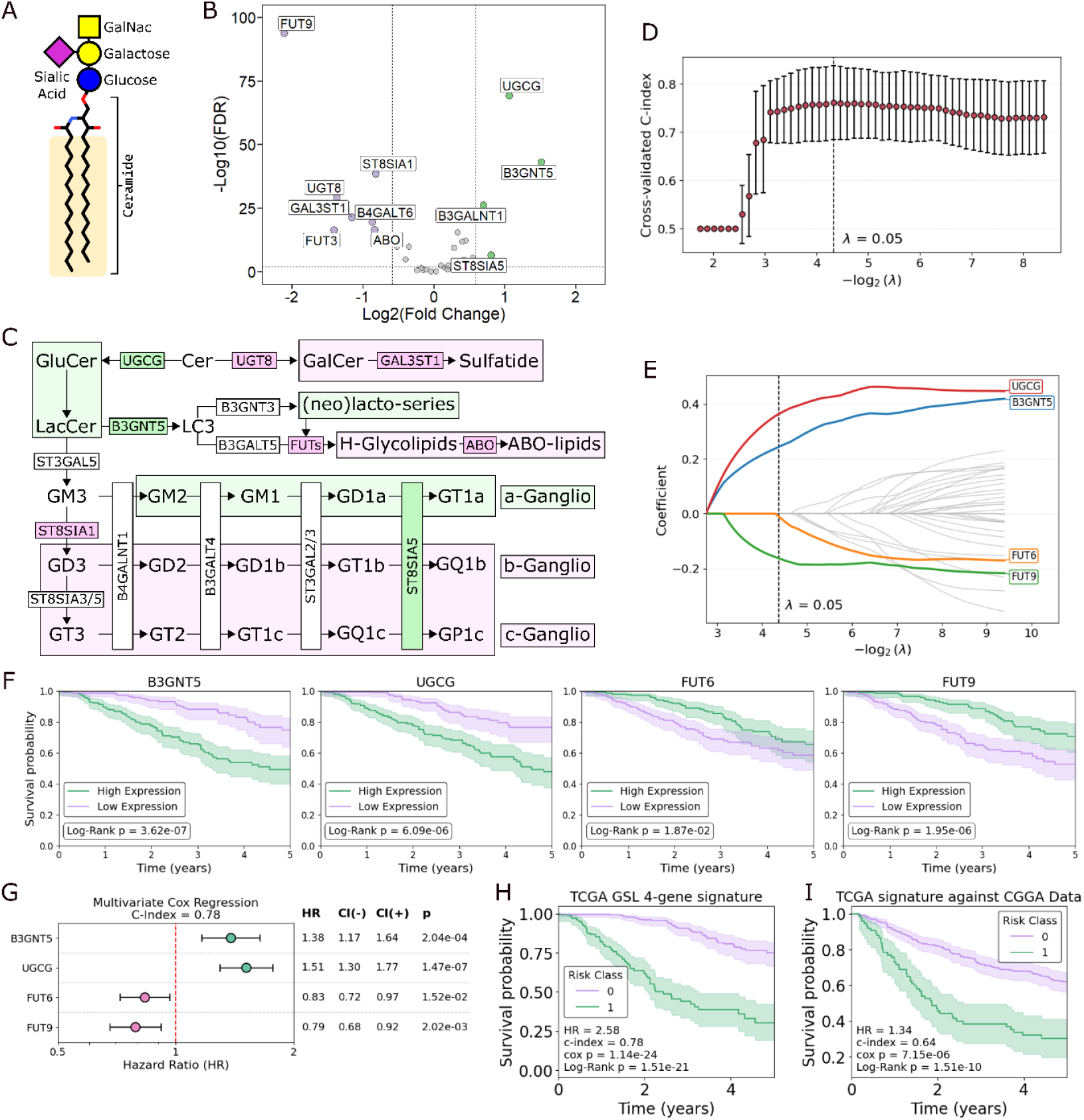
Analysis of glycosphingolipid (GSL) differential gene expression in Glioblastoma Multiforme (GBM) and Lower Grade Glioma (LGG). **A**. Schematic diagram of a model GSL, the ganglioside GM2. The ceramide tail and individual head group sugars are labelled. **B**. Differential expression analysis of TCGA RNA-seq data for GBM against LGG. Significantly upregulated genes in GBM are highlighted in green, whilst significantly downregulated genes are highlighted in purple. A >1.5-fold change and *p* <0.01 thresholds were used to determine significance. **C**. Schematic diagram of part of the GSL synthetic pathway, highlighting changes in gene expression in GBM vs LGG in corresponding colours. Potentially impacted lipids/lipid-series are also highlighted to align with the differential expression data. **D**. Cross-validated Lasso regression against expression values for 34 genes of glycosphingolipid synthesis from TCGA LGG records, scoring against Cox proportional hazards multivariate c-index. The optimum λ parameter (maximising c-index) is highlighted by a dotted line. **E**. Cox coefficients from multivariate Cox regression experiments against TCGA gene expression data for LGG patients. The optimal lambda of 0.05 is shown with a dotted line, with non-zero coefficient genes at this λ highlighted in colour. **F**. Kaplan-Meier (KM) analysis of the four genes identified in panel E. High and low expression groups were determined via stratification around the median expression value for each gene, using TCGA LGG data. **G**. Multivariate Cox regression of TCGA LGG data performed against the 4 selected genes, showing hazard ratios in the plot window, with 1.0 marked as a red line. **H**. KM analysis of TCGA risk stratification on the GSL risk score with a log-rank test tuned decision threshold. **I**. KM analysis of CGGA LGG data stratified by a risk score derived from the multivariate Cox regression coefficients for each of the 4 genes in panel G, based on TCGA data.

GSLs are synthesised in the endoplasmic reticulum and Golgi apparatus via a series of stepwise additions of different glycan moieties. This glycan headgroup synthesis involves several catalytic enzymes including glycosyltransferases, sialyltransferases and fucosyltransferases[24–26]. The combined gene expression and activity of these enzymes produce the different GSL repertoires present in a given cell type. This GSL repertoire changes during development and is cell-type specific[27]. The restricted expression of some GSL subtypes, such as GD2 and fucosylated GSLs, has made them ideal for targeted therapies[15]. Furthermore, inhibition of GSL synthesis can impair tumour growth and induce apoptosis[28].

Although individual GSL synthetic enzymes have been associated with specific cancers, there has been very limited exploration of how the broader family of synthetic genes are expressed in cancers and there has been no incorporation of such data into metabolic models for prediction of altered overall GSL repertoire. Advancements in machine learning strategies and metabolic modelling pipelines, combined with publicly available, highly curated transcriptomics datasets are enabling new discoveries in cancer research[29]. Here we explore the role of GSL synthetic enzyme gene expression in LGGs and identify that specific changes in three key genes are highly predictive for poor LGG prognosis. This identification of the potential role of GSL changes in the transition from LGG to GBM will help contribute to improved patient prognostic stratification to optimise clinical treatment strategies[4]. Furthermore, the known functional roles of GSLs in altered cell signalling may inform potential future therapeutic avenues targeting GSL-modulated pathways or GSL synthesis directly.

## Results & Discussion

### GSL genes are prognostic indicators in LGG as well as being upregulated in GBM

The first approach taken to investigate GSL metabolism as a candidate for risk modelling in LGG was to perform differential expression analysis between the gene expression profiles of LGG patients and the more severe GBM patients in The Cancer Genome Atlas (TCGA). Eleven key genes of glycosphingolipid biosynthesis were significantly differentially expressed between the two cancers (p < 0.01, >1.5 FC). Genes that were downregulated in GBM compared to LGG were the fucosyltransferases FUT9 and FUT3, the Galactosylceramide (GalCer) synthase UGT8, the sulfatide synthase GAL3ST1, one of the LacCer synthases B4GALT6, histo-blood group ABO system transferase ABO and the sialyltransferase ST8SIA1. Those that were instead upregulated were the LC3 synthase/(neo)lacto-series gatekeeper B3GNT5, the Glucosylceramide (GluCer) synthase UGCG, the sialyltransferase ST8SIA5 and the globotriaosylceramide synthase B3GALNT1 (**Fig.1B**).

To better interpret the potential implications of these expression changes, they were mapped onto a reduced schema of GSL metabolism (**Fig.1C**). The core unit of a GSL is the ceramide backbone which can be initially modified by the addition of either a glucose via UGCG or alternatively a galactose via UGT8. This represents the first major metabolic bifurcation in GSL synthesis. The observed downregulation of UGT8 combined with the upregulation of UGCG, suggests that the most apparent of the expression-inferred changes to GSL metabolism would be an expected shift towards GluCer-based lipids, away from GalCer-based lipids in GBM when compared to LGG.

Downstream of GluCer, there are further gene expression changes in GBM that have the potential to redirect GSL metabolism to alter the overall GSL repertoire of GBM versus LGG. The increase in B3GNT5 expression, alongside the decrease in fucosyltransferase (FUT9/FUT3) and ABO expression could suggest a prioritisation of (neo)lacto series lipids over the H- and ABO-glycolipids. Weaker inferences can be made for the gangliosides, where decreased ST8SIA1 expression, coupled to increased ST8SIA5 expression could imply a push towards the more complex a-series gangliosides.

Following the differential expression analysis, to identify LGG-specific risk factors, Cox regression and Kaplan-Meier (KM) analyses were performed on TCGA LGG data. Elastic net lasso regression was tuned under cross-validation, identifying λ = 0.5 to maximise c-index (**Fig.1D**). The subsequent Cox regression identified 4 GSL synthetic genes with non-zero coefficients at λ = 0.5 out of the 34 genes included in the analysis. These genes were UGCG, B3GNT5, FUT6 and FUT9 (**Fig.1E**). Interestingly, the enzymes with positive coefficients, UGCG and B3GNT5 in the LGG analysis, are also upregulated in GBM when compared to LGG, whilst the FUT9s negative coefficient aligns with its downregulation in GBM. This observation, that the genes that are implicated in poor prognosis within LGG are the same as those that are upregulated in more severe GBM strongly suggests a role for GSL metabolism in disease severity and potentially progression from LGG to GBM. KM curves for the 4 identified genes are shown in **Fig.1F**, with risk groups separated around median expression.

To generate a 4-gene risk signature, multivariate Cox regression was performed on TCGA data (**Fig.1G**). The 4-gene score was defined by the coefficient-weighted sum of expression values for each gene (Equation 1). The TCGA-trained coefficients were used to calculate risk scores for LGG data from a completely independent validation set, the Chinese Glioma Genome Atlas (CGGA). The risk group threshold was this time tuned on TCGA data based on maximising the log-rank test statistic. Resulting KM analysis showed significant risk group stratification against the CGGA data (Log-Rank p = 1.5e-10). Univariate Cox analysis of the continuous risk score yielded a significant result against both the TCGA data and the CGGA validation data, with hazard ratios of 2.58 (*p* = 1.1x10^-24^, **Fig.1H**) and 1.34 (p=7.2x10^-6^, **Fig.1I**). This result is comparable to similar analysis for LGG previously performed on fatty acid metabolism against TCGA data, yielding a 4-gene risk signature with a hazard ratio of 1.57 (p <0.01)[30].

### Computational modelling of GSL metabolism with pyGSLModel as a feature engineering strategy for survival prediction

The GSL repertoire of a cell is not solely determined by the level of gene expression of the synthetic enzymes. The availability of substrates and enzyme-substrate preferences will contribute to the final GSL composition. Changes in gatekeeper/branchpoint enzyme expression, such as UGCG, UGT8 and B3GNT5, might be expected to result in significant shifts in the GSL composition of a cell, but multiple smaller changes may also have the potential to shift GSL repertoires in complex and difficult to predict ways. With this in mind, despite the promising performance of the 4-gene GSL signature for survival modelling, metabolic simulation was utilised as a feature engineering approach to buttress model training.

We developed a python package (pyGSLModel) to implement constraint-based metabolic simulations built on top of COBRApy[31], pyFastCore[32] and iMATpy[33]. The iMAT method was used to perform patient-specific simulations of GSL metabolism based on their GSL synthetic enzyme expression data using a small-scale metabolic model of GSL synthesis pruned from the HUMAN-GEM model[34] in pyGSLModel, preserving core reactions of metabolism and reactions of GSL synthesis. This approach enabled the incorporation of relationship information between GSL synthetic genes based on their relative positions within the metabolic network, with this information, in principle, represented in the corresponding reaction fluxes for a patient.

Transcriptomic integrative metabolic simulations were performed for TCGA and CGGA LGG data for genes of GSL synthesis across 24 different sets of iMAT input parameters (Table 1). Considerable variation in output can be seen across iMAT parameters (**Fig.2**), demonstrating the need to trial different parameters and evaluate their impacts on model performance empirically. Generally, a shift away from globo-, (neo)lacto- and ganglio c-series towards gal-series and a/b-series gangliosides was observed when increasing the epsilon (the minimum flux required for a reaction to be considered ‘active’) and threshold (the flux limit above which a lowly expressed reaction is penalised for being ‘active’) values. These two parameters both contribute to the objective function for the simulation which seeks to maximise the consistency between the data and the model. This parameter sensitivity highlights the need for caution when trying to make biological inferences based on these kinds of metabolic simulation, despite their potential empirical benefit for modelling tasks.

**Table 1.**
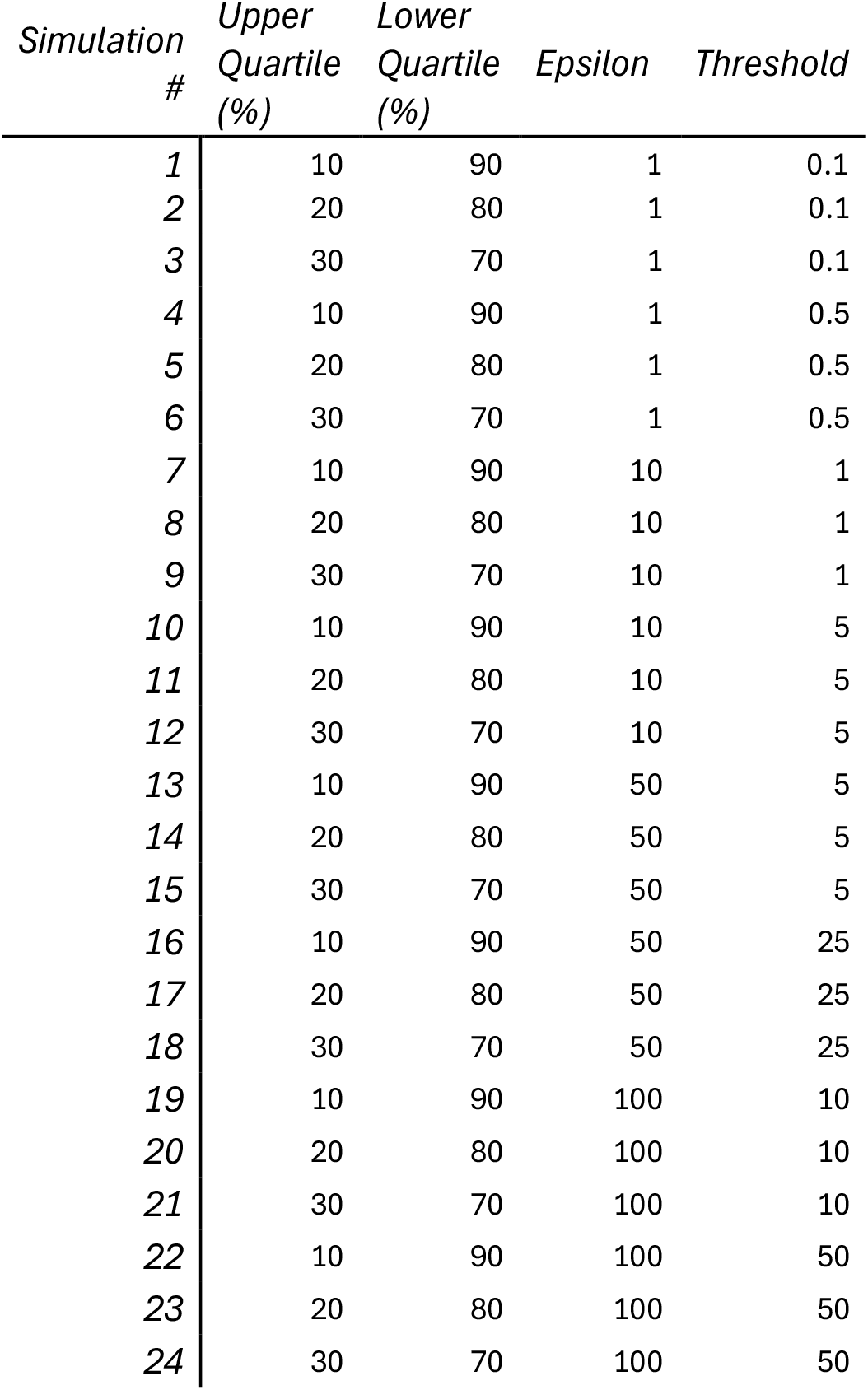
Set of parameters trialled for iMAT integration.

**Figure 2.**
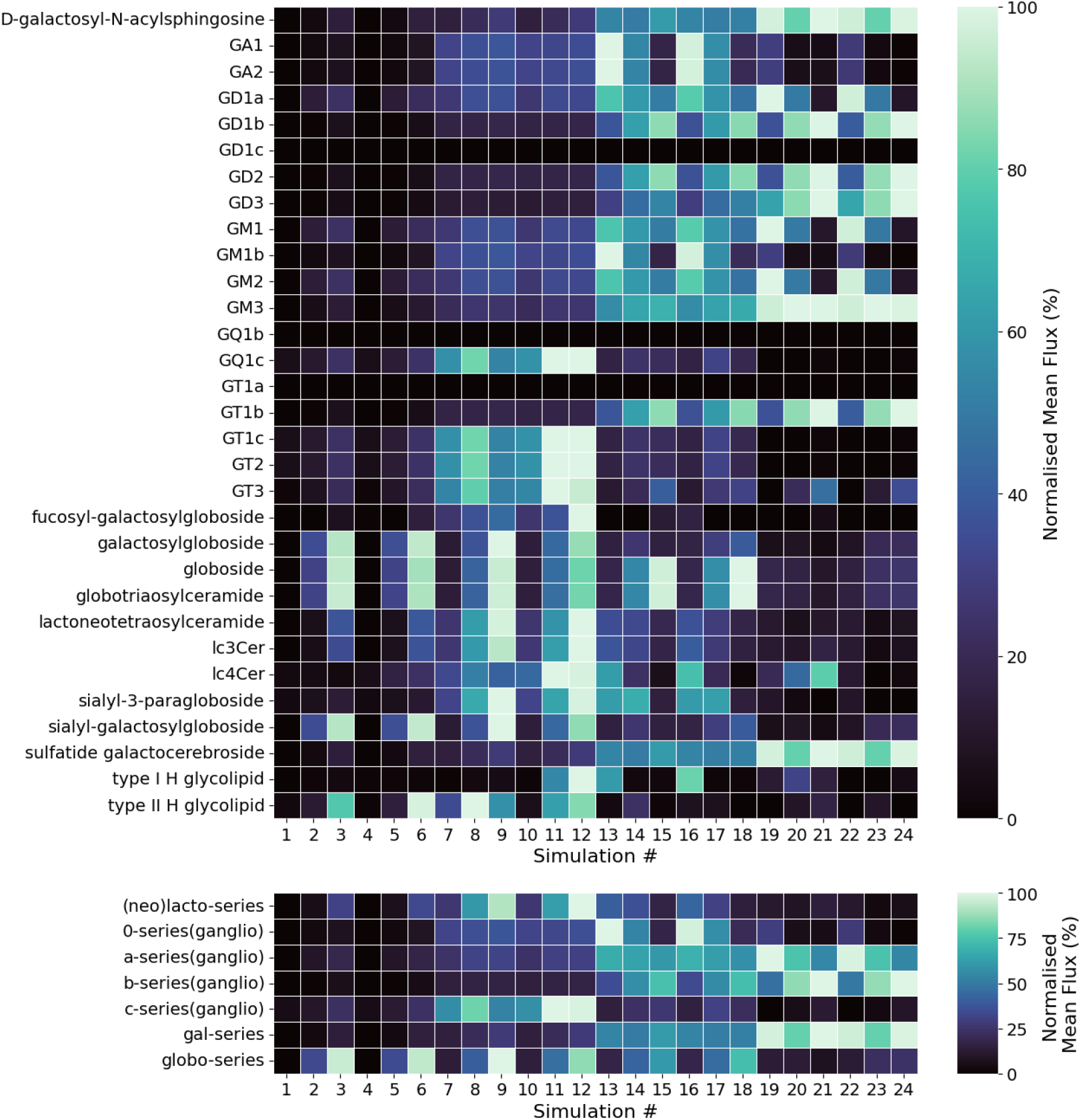
GSL metabolic flux analysis. Normalised mean simulated fluxes across all TCGA LGG patients across each set of simulation parameters (Table 1). Simulated fluxes across series were summed to give the total Normalised Mean Flux per lipid series.

Separate datasets were prepared corresponding to each different parameter set for model training, in which the simulated metabolic fluxes for GSL synthetic reactions were appended onto GSL-synthetic enzyme expression data. This resulted in 24 separate datasets to be trialled in subsequent model development (**Fig.3**).

**Figure 3.**
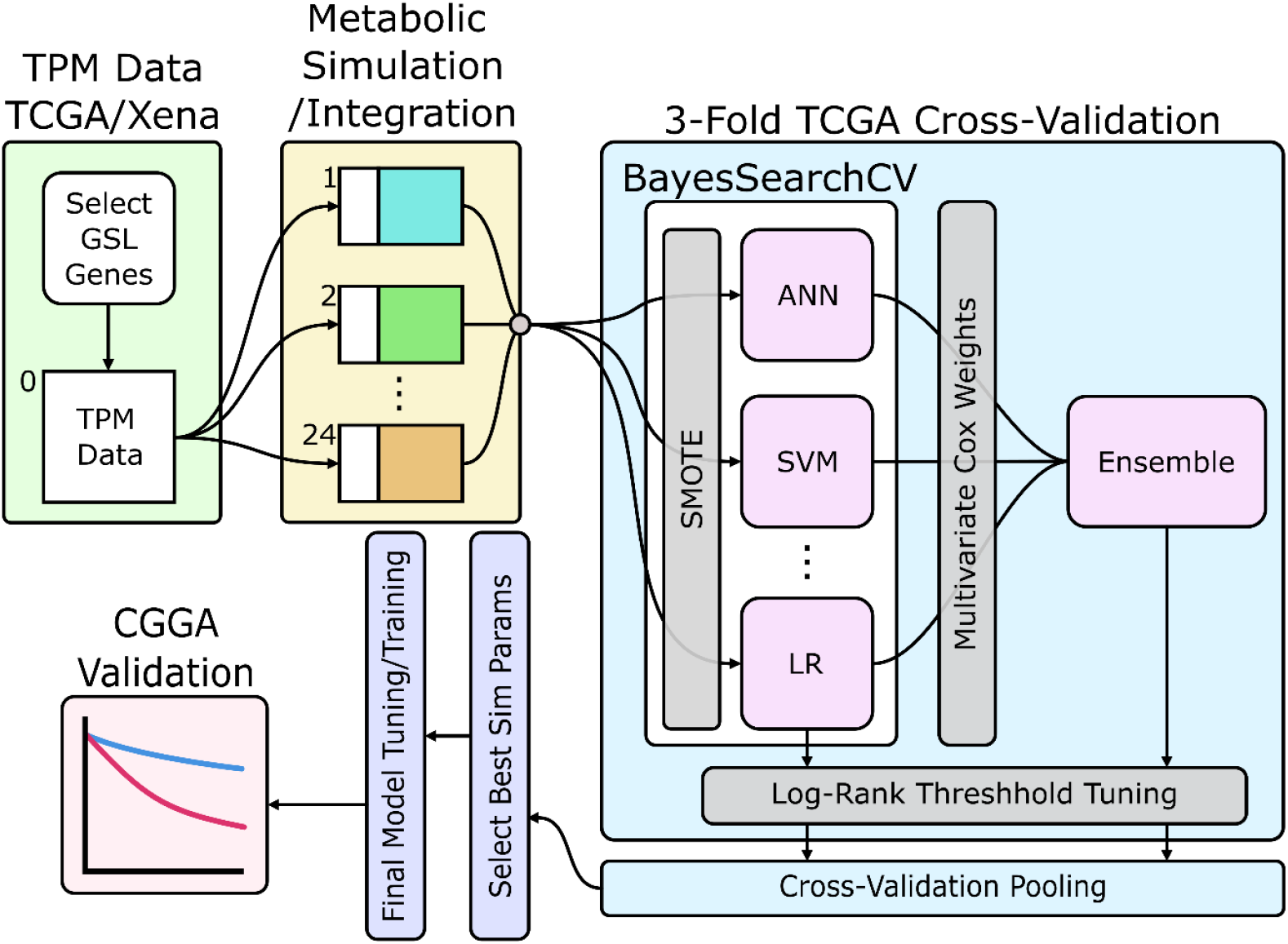
Schematic of the workflow of this study. Green box shows GSL synthetic gene TPM expression data accession from the TCGA via Xena. Yellow box shows the integration of simulated metabolic data to append to the expression dataset. Light Blue box shows the nested cross-validation (CV) machine learning approach. Data underwent synthetic minority oversampling (SMOTE) to deal with class imbalance before training 5 different models to predict overall survival: Artificial Neural network (ANN), Support Vector Machine (SVM), RandomForest (RF), XGBoost (XGB) and Logistic Regression (LR). Model output probabilities were then combined to generate an ensemble model based on multivariate Cox coefficients for each model. Threshold tuning was performed and cross-validation results pooled for analysis. Dark blur box shows final model training and cross-validated tuning trained on all TCGA data, using metabolic simulation parameters identified in the nested CV experiment. Pink box denotes final model validation against completely independent LGG data from the CGGA.

### Ensemble machine learning approaches effectively model survival based on GSL synthesis

To generate a GSL risk score that captures patterns of change across GSL synthesis, rather than traditional gene signatures such as was demonstrated above (**Fig.1H**), an ensemble machine learning approach was implemented. This method allows us to utilise the full GSL synthetic gene expression data available, alongside the appended metabolic simulation reaction fluxes. An ensemble approach was chosen, rather than focusing on any one specific model to offer increased robustness in final score calculation and risk stratification, minimising reliance on a small number of signals that a single model may become overly reliant on. Five different machine learning models were trained, a Support Vector Machine (SVM), a RandomForest (RF) model, an XGBoost (XGB) model, a Logistic Regression (LR) model and an Artificial Neural Network (ANN). Each model was trained separately on each of the 24 prepared datasets for each set of simulation parameters to predict overall survival (OS). SMOTE oversampling was used to address class imbalance. The models were trained under nested cross-validation and Bayesian optimisation against solely TCGA data in the first instance. The ensemble model was engineered by performing multivariate Cox regression against the 5 trained models and subsequently weighting their probabilities by the Cox coefficients before calculating the weighted sum (Equation 2). This approach enables a survival-specific method for model weighting to produce the ensemble score. The resulting models were evaluated against the pooled cross-validation sets. Model thresholds were tuned based on identifying probability/ensemble score thresholds that maximised the log-rank test statistic. This approach is outlined in **Fig.3**.

This process was repeated 5 times at different random states. The best performing set of simulation parameters (in terms of c-index) across repeats was selected to take forward for final model training. iMAT simulation parameters of: upper quartile=30%, lower quartile=70%, ε=100, threshold=10 achieved the best performance, with an average c-index of 0.77. This corresponded to parameter set 21.

The final models were trained using all TCGA data and underwent cross-validated Bayesian optimisation. The resulting model was validated against the independent CGGA dataset. Cox coefficients for ensemble weighting and the risk classification threshold were defined using only the training TCGA data to avoid data leakage. Identical genes of GSL synthesis were used and underwent metabolic simulations as per parameter set 21.

KM analysis of predicted risk groups showed significant stratification across all 6 models with the high-risk groups showing a steep decline in survival at < 2 years to ≈ 0.2 survival probability. From 2 to 5 years, the rate of decrease in survival probability levelled off as most high-risk patients died in the 0–2-year time window. In contrast to this, the low-risk group showed an approximately linear decrease in survival over the 5-year window from ≈ 1.0 to 0.6 (**Fig.4A**). The distribution of true deaths along the ensemble score (**Fig.4B**) demonstrates exceptional stratification around the tuned threshold, and that the selection of a more forgiving threshold would have resulted in considerable contamination from false positives.

**Figure 4.**
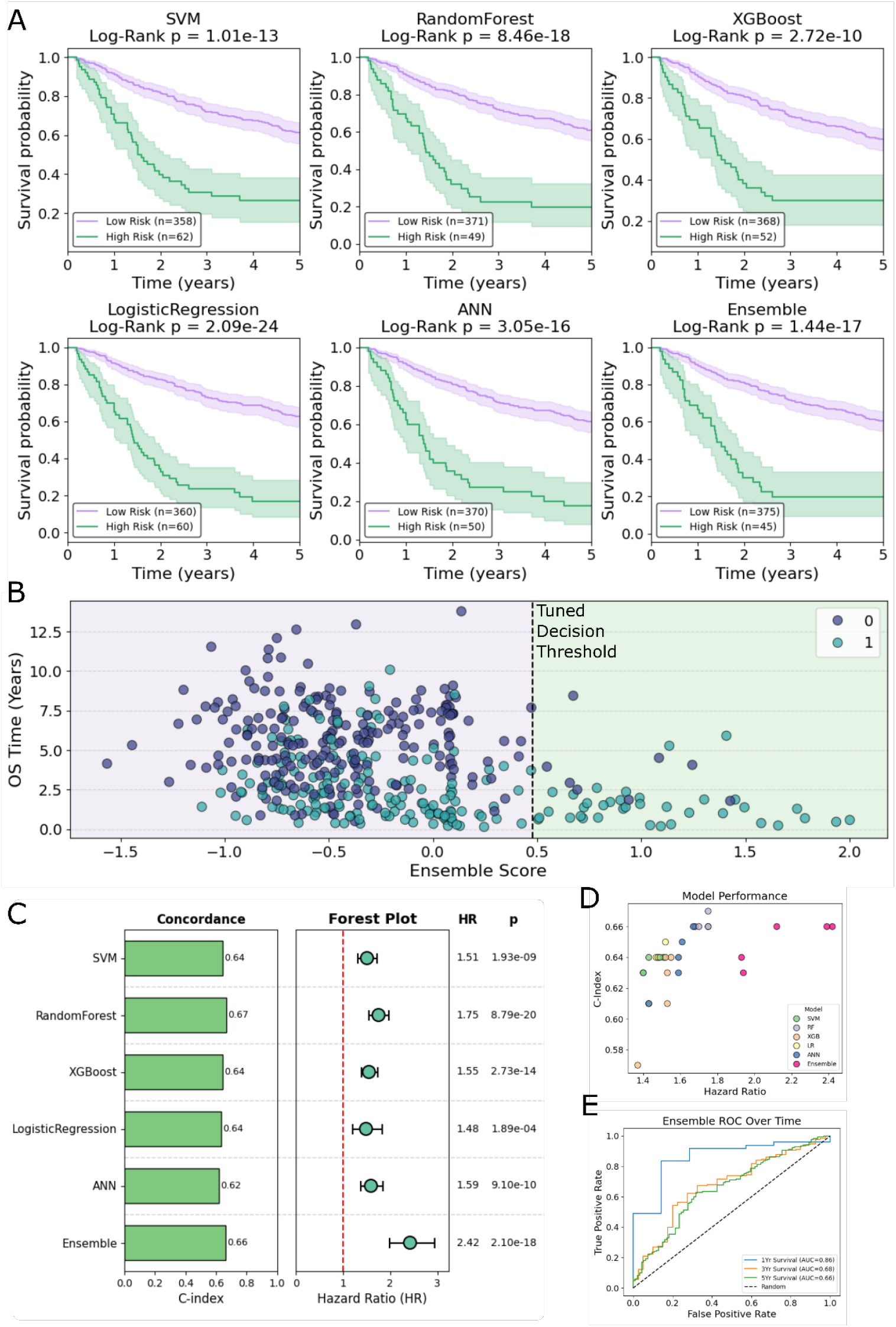
Patient stratification with ensemble machine learning. **A** KM survival curves showing stratification between ML model defined risk groups for CGGA data (models trained on TCGA data exclusively). Low-risk predictions are in purple, whilst high-risk are in green. **B**. Overall Survival (OS) time plotted against ensemble model scores. **C**. Output performance statistics for each ML approach for the representative random state. **D**. Model performance plot of c-index against hazard ratio. Each random state replicate for each model is plotted with colours separating the different ML approaches. **E**. Receiver-operator-characteristic curve for a representative ensemble model (consistent with panel A). Data is separated by time intervals, showing curves for <1 Yr, <3Yr and <5yr survival. For panels **A, B, C & E**, a representative random state (consistent across panels) is used to represent results.

Each model’s continuous probability/score outputs were evaluated with univariate Cox regression. All models showed significant results, with the ensemble model performing particularly well with a c-index of 0.66 alongside a hazard ratio of 2.42 (**Fig.4C&D**). This result improves against the traditional multivariate Cox approach, which, when validated against CGGA data, achieved a c-index of 0.64 and hazard ratio of 1.34 (**Fig.1H**). Examining the ROC curve over time for the ensemble model (**Fig.4E**) demonstrates that the model’s best performance occurs when predicting particularly severe cases with < 1 year survival. In this case, the Ensemble model shows an ROC-AUC of 0.86 compared to 0.68 and 0.66 for <3 year and <5 year survival, respectively.

### Adhesion, motility and immune infiltration are enriched in GSL-score high-risk groups

To exploit the biological relevance of the model outputs, we performed differential expression analysis between high- and low-risk groups predicted by the Ensemble model. We did this for each of 5 repeat experiments for the TCGA model training and CGGA validation, performed at different random states. Genes that were significantly upregulated in every repeat were selected for subsequent gene ontology analysis via DAVID[35]. 1079 genes were identified and analysed for enrichment across GO Direct Biological Processes (BP), Cellular Component (CC) and Molecular Function (MF). Functional annotation clustering was performed[36] to identify more specific biological insights (**Fig.5A**).

**Figure 5.**
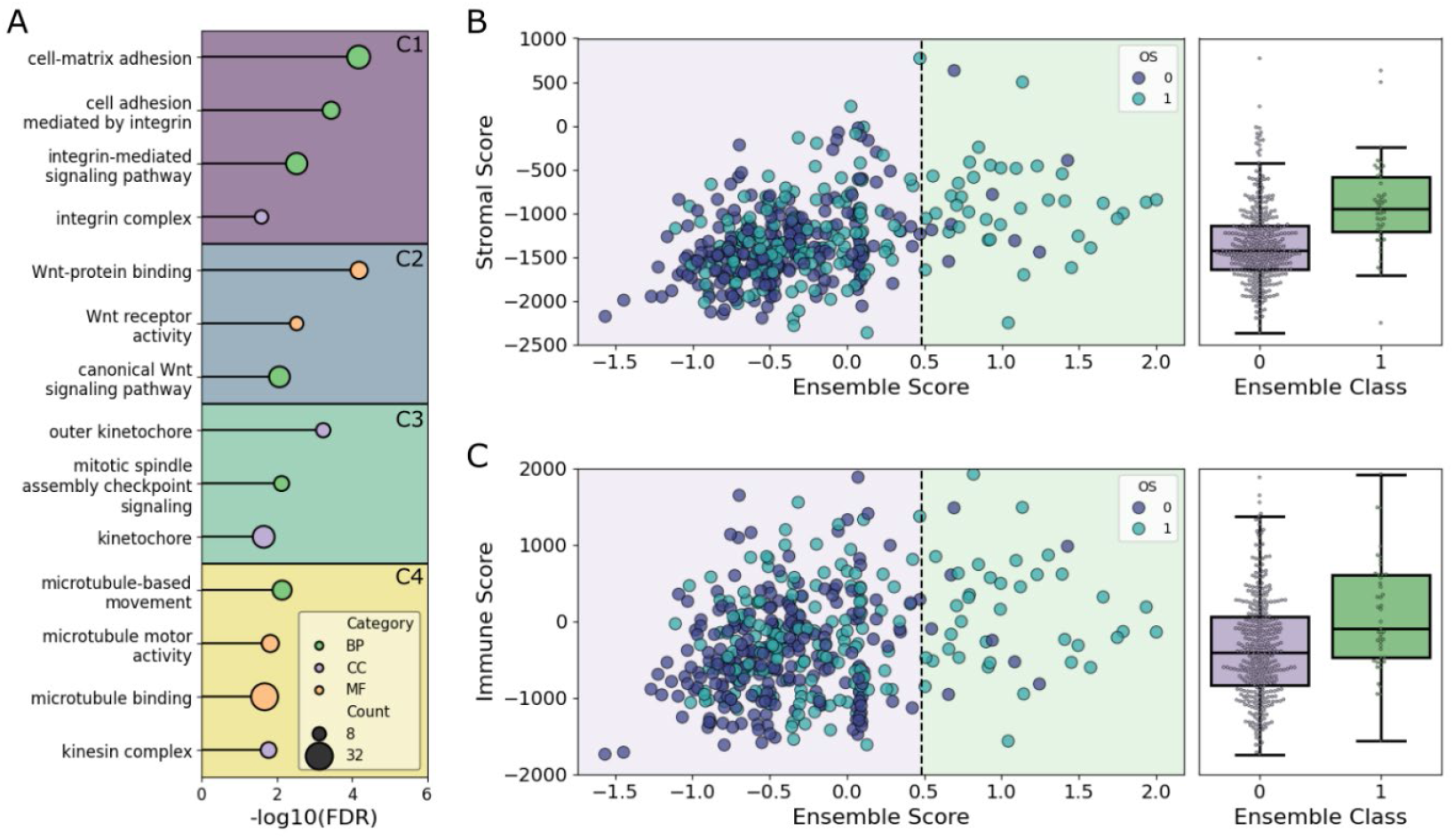
Functional analysis of model derived risk groups. **A** Functional annotation clustering with DAVID[36] of consistently upregulated genes in the predicted high-risk group for the ensemble model across all repeats. Clusters are labelled C1-4 and separated by coloured boxes. GO Direct terms are denoted by stem plot point colours. Biological Process (BP) are shown in green, Cellular Compartment (CC) in purple and Molecular Function (MF) in orange. **B** Left panel: Stromal score calculated via the Estimate algorithm[51] against Ensemble model scores (for a representative model). Decision threshold denoted by the dotted black line, with the low-risk region highlighted in purple, and high-risk in green. Points are coloured by overall survival, with death represented by light blue, and unrecorded as dark blue. Right panel: Boxplots and points showing stromal score (linked axis to left panel) against Ensemble predicted class. **C** Left panel: Immune score calculated with the Estimate algorithm[51] against Ensemble model scores (for a representative model, consistent with panel B. Data presented as in panel B.

The most enriched cluster, with a cluster enrichment score of 5.12 included four GO terms relating to integrin signalling and cell adhesion. This observation is consistent with the well-characterised association between GSL synthesis and cell adhesion particularly the detrimental impact of gangliosides loss in the central nervous system[37,38]. In addition to reduced cell adhesion, disruption of GSL biosynthesis can also impact integrin gene expression[39]. Importantly, the specific shift between the production of sulphated versus sialylated GSLs has differential effects on integrin and growth factor receptor gene expression. Recent work has also identified a growth factor-triggered GSL-driven mechanism by which integrins are removed from the plasma membrane surface to mediate cell motility[40]. Together, these data suggest a direct mechanistic link between the GSL composition of a cell and integrin-mediated cell adhesion.

Three Wnt signalling associated genes also formed an enriched cluster (cluster enrichment score = 4.56). The activation of Wnt signalling involves the binding of Wnt proteins to their receptor complexes in ordered plasma membrane microdomains enriched with sphingolipids[41]. LRP6 is an essential co-receptor for Wnt signalling and binds the GSL GM1 within membrane microdomains[42]. Disruption of membrane order and altered GSL metabolism interfere with the phosphorylation of LRP6 disrupting downstream signalling[41]. Interestingly, the GSL mannosyl glucosylceramide (MacCer) promotes synaptic bouton assembly at the neuromuscular junction in *Drosophila* by binding and enhancing the levels of Wnt1/Wingless at the plasma membrane, facilitating presynaptic Wnt signalling[43]. Misregulation of Wnt signalling is linked to disrupted plasma membrane lipid organisation in a range of cancers, with Wnt signalling being one of the key processes in the regulation of cell stemness, which is often coopted in cancer[44,45]. This aligns with previous literature highlighting that B3GNT5 gene expression increased the stem-like behaviour of cells in GBM[22]. Additional studies have further implicated B3GNT5 in tumour promotion and stemness of a range of cancers[23,46].

Kinetochore and mitotic spindle associated GO terms were identified in cluster 3, with an enrichment score of 4.10, suggesting implications in cell division for the GSL risk group. Several enzymes involved in GSL metabolism have been shown to play important roles in cytokinesis, including GAL3ST1, ST8SIA5 and UGCG[47,48]. Individual depletion of expression of these genes results in failure of cell division potentially due to the mislocalisation of cytoskeletal proteins that connect the plasma membrane to the actin cortex. In addition to mislocalisation, altered GSL expression changes the expression of cytoskeletal genes. The depletion of UGCG gene expression reduced stem cell proliferation and decreased the expression of microtubule-associated protein 2 (MAP2) in neurons and glial fibrillary acidic protein (GFAP) in glial cells[49]. Potentially related to this, the fourth GO term cluster (with an enrichment score of 3.28) relates to microtubule movement, binding and the kinesin complex. Sphingolipid-rich membrane microdomains dynamically interact with the underlying cytoskeleton regulating signalling and mediating communication across the plasma membrane[50]. Although the molecular details of this process remain poorly understood, the impact of altered GSL metabolism on the cytoskeleton in brain cancers may provide an exciting area of research.

Immune and stromal infiltration scores were calculated via the Estimate algorithm[51] and compared across GSL risk-scores and risk groups (**Fig.5B&C**). For both immune and stromal scores there is a subtle increase in high GSL-score patients. This, alongside the array of immune-related GSL functions[18], marks GSL-related immune dysfunction at tumour sites another potential point of study for therapeutic potential in glioma patients[28].

### The potential for GSL-based prognostics

The effective risk stratification achieved through our ensemble machine learning approach suggests the expression and metabolic network of GSL synthetic genes carries information highly relevant to disease severity in LGG. It can be postulated that in these high-risk cases predicted with our GSL based method there will be dysregulated GSL synthesis and correspondingly aberrated GSL enrichment in cells. Considering the shedding of GSLs into the tumour microenvironment, with their detection in patient serum demonstrated in a number of cancers[52], GSL based prognostics and diagnostics may offer a useful tool to identify high-risk patients. Depending on the overall downstream consequences to tumour aggression of disrupted GSL synthesis, such high-risk patients may benefit from treatment supplementation with GSL targeting therapies. These could both act to inhibit GSL synthesis or also exploit GSLs to target the cancer. Such approaches have already been exploited for Neuroblastoma, where the monoclonal antibody Dinutuximab binds GM2 to target the cancer cells[53–55] Tumour-associated-gangliosides have also shown promise for early-stage detection in ovarian cancer[56].

With these models being trained on limited, open-source data, the potential for further development with more extensive clinical data is considerable. This is particularly relevant as society moves towards precision medicine approaches, where these types of methods could provide highly specific treatment strategies on a patient-by-patient basis. This GSL specific approach could easily be multiplexed alongside equivalent methods focusing on different metabolic pathways, giving patients biologically relevant risk stratification to inform clinical decision making.

Furthermore, the clear relevance of GSL biology demonstrated here to patient outcomes in LGG suggests new avenues for investigation to understand mechanisms driving disease severity.

## Methods

### Data Accession

Training data for LGG (grades III and IV) were accessed from The Cancer Genome Atlas (TCGA) via the Xena Database[57]. Transcripts per Million (TPM) gene expression data for GSL synthetic enzymes were acquired from the TCGA-TARGET-GTEx dataset hosted on Xena[57,58].

Raw count data and metadata for both LGG and GBM were also accessed from the TCGA via the Xena database for differential expression analysis.

Validation data were accessed from the Chinese Glioma Genome Atlas (CGGA)[59] as FPKM values which were converted to TPM values and Log2(x+0.01) normalised before filtering to the relevant GSL synthetic genes to match the training data format.

### Differential Expression Analysis

All differential expression analyses were performed in R using the DESEQ2 package[60].

Raw count data was used, with low count genes being filtered out before analysis. Complete scripts for the differential expression analysis are available on Github (https://github.com/JackWJW/LGG_Prognosis_Prediction).

### Survival Analysis

Kaplan-Meier (KM) analysis was conducted in the lifelines package in python[61]. Elastic Net regression analyses were performed with the scikit-survival package in python[62]. Full lasso regression was performed with scikit survival L1 regularisation set at 0.5.

Cox regression analysis of GSL genes from the TCGA data was performed with the lifelines package in python, utilising the best alpha (0.05) identified from lasso/elastic net regression (0.5 L1 regularisation) experiments. 0.05 was utilised as the L2 regularisation for subsequent ridge Cox regression analyses performed against genes and models.

The 4-gene-signature was calculated by performing multivariate regression against the 4 key genes identified from the Lasso regression feature selection experiment and subsequently weighting expression values for each gene by their Cox coefficients to calculate their aggregate risk score (Equation 1).

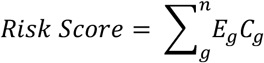

Where *E*_*g*_ refers to a genes expression and *C*_*g*_ refers to a genes Cox coefficient.

### Metabolic Modelling and Simulations

Metabolic modelling was performed using the custom written pyGSLModel python package. pyGSLModel was built on top of COBRApy[31], iMATpy[33] (https://pypi.org/project/imatpy/) and pyfastcore[32] (https://pypi.org/project/pyfastcore/) to simplify the preparation of sphingolipid metabolism specific models and subsequent transcriptomic integration and metabolic simulation. A sphingolipid specific metabolic model was derived from the human genome scale HUMAN-GEM model[34]. Trial simulations were run on the HUMAN-GEM with a neuronal GSL-specific objective function defined based on key GSL proportions known in neuronal cell models[63]. Reactions carrying flux in trial simulations as well as manually curated reactions affiliated with sphingolipid catabolism were defined as core reactions before performing automated model pruning of the HUMAN-GEM with pyGSLModel utilising pyfastcore. This process resulted into a specific model of glycosphingolipid synthesis.

Transcriptomic integration was performed via the iMAT method which was implemented via iMATpy in pyGSLModel. With respect to training TCGA data, iMAT simulations were performed for each LGG sample in the dataset leveraging 34 genes of GSL synthesis to define iMAT rankings. The pruned metabolic model of GSL synthesis was utilised for all iMAT simulations. To enable the identification of the best performing simulation parameters, 24 different parameter sets were trialled. Each parameter set includes variable values for the upper quartile (% of highest expressing genes to be denoted +1 for iMAT), lower quartile (% of lowest expressing genes to be denoted -1 for iMAT), threshold (flux above which the solution score is increased for +1 reactions) and epsilon (flux above which the solution score is penalised for -1 reactions). Parameter sets are shown in Table 1.

CGGA data for GSL synthetic genes underwent iMAT simulation with parameters from the best performing parameter set (identified in TCGA cross-validation).

pyGSLModel iMAT integration functions also calculate total fluxes throughout lipid-series as well as individual fluxes for reactions of GSL synthesis. These total lipid-series fluxes were also included in the simulated flux appended datasets for model training and validation.

### TCGA Model Cross-Validation

TCGA-only models were trained under nested cross-validation with 3 stratified outer and inner folds. In the data preparation pipeline data underwent standard scaling (sklearn) alongside SMOTE oversampling (imblearn). All models underwent 25 iterations of Bayesian optimisation (BayesSearchCV, sklearn) in the inner CV loop, with scoring on average precision (area under the precision-recall curve). Outer fold cross-validation results were pooled for evaluation. This process was repeated for each different set of iMAT parameters, at 5 different random states (0-4). SVM (sklearn), RF (sklearn), XGB (XGBoost), LR (sklearn) and ANN (pytorch, skorch) models were trained with this approach. The ANN was trained for a maximum of 300 epochs with early stopping, focal loss and the Adam optimiser[64–67]. The ensemble prediction was generated by performing multivariate Cox regression of the 5 model’s probabilities for validation samples against overall survival and survival time. The resulting Cox coefficients were utilised as weights for ensemble model calculation (Equation 2).

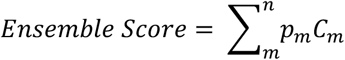

Where *p*_*m*_ refers to model probability and *C*_*m*_ refers to model Cox coefficient.

### Final Model Training (TCGA Data) and Evaluation (CGGA Data)

The final model was trained on all TCGA data with Bayesian optimisation performed under 5-fold cross-validation for 50 iterations with scoring on average precision. In the data preparation pipeline, data underwent standard scaling (sklearn) alongside SMOTE oversampling (imblearn). This process was repeated across 5 different random states (0-4). SVM, RF, XGV, LR and ANN models were trained with this approach. The ANN was trained for a maximum of 300 epochs with early stopping, focal loss and the Adam optimiser. The ensemble model generated its prediction as previously, via multivariate Cox regression of model probabilities to generate Cox coefficients to calculate a Cox weighted ensemble score (Equation 2).

The resulting models were validated against CGGA data.

### GO Analysis

Differential expression analysis was performed between high-risk and low-risk groups predicted by the ensemble model against CGGA data. This was done for each of the 5 repeats, selecting genes that were consistently upregulated in all repeats to take forward for Gene ontology (GO) analysis.

Gene ontology (GO) analysis was performed using the DAVID web server[36] against GO Direct terms for Biological Processes (BP), Cellular Compartment (CC), and Molecular Function (MF). Functional annotation clustering was performed on the “Medium” Classification Stringency with an EASE Threshold of 1. Results from clustering were filtered for cluster enrichment scores > 3.0 and Benjamini-Hochberg adjusted p-value <= 0.05.

### Tumour Purity Analysis

Tumour purity, immune, stromal and estimate scores were calculated using the estimate algorithm (tidyestimate package in R) against CGGA FPKM data for all genes and samples.

### Data Visualisation

Differential expression volcano plots were generated with the ggplot package in R[68]. All other data visualisation was performed using the matplotlib and seaborn packages in python[69,70].

## Data Availability

All data, code and models are available on Github (https://github.com/JackWJW/LGG_Prognosis_Prediction) and Huggingface (https://huggingface.co/JackWJW/LGG_Prognosis_Ensemble). The pyGSLModel python package is available on pypi (https://pypi.org/project/pyGSLModel/), with the complete code on Github (https://github.com/JackWJW/pyGSLModel), providing functionality for GSL-specific metabolic simulations and calculating GSL-risk scores from RNA-seq data.

## Acknowledgements

We thank Daniel Machado for helpful early discussions regarding metabolic modelling strategies. This work was supported by a Wellcome Trust Senior Research Fellowship (219447/Z/19/Z) to JED. JWJW is supported by a Michael House PhD studentship. The results published here are in part based upon data generated by the TCGA Research Network: https://www.cancer.gov/tcga. For the purpose of open access, the author has applied a Creative Commons Attribution (CC BY) license to any Author Accepted Manuscript version arising from this submission.

